# Equilibrium dialysis with HPLC detection to measure substrate binding affinity of a non-heme iron halogenase

**DOI:** 10.1101/2024.04.03.588023

**Authors:** Elizabeth R. Smithwick, Ambika Bhagi-Damodaran, Anoop Rama Damodaran

## Abstract

Determination of substrate binding affinity (*K*_d_) is critical to understanding enzyme function. An extensive number of methods have been developed and employed to study ligand/substrate binding, but the best approach depends greatly on the substrate and the enzyme in question. Below we describe how to measure the *K*_d_ of BesD, a non-heme iron halogenase, for its native substrate lysine using equilibrium dialysis with subsequent detection with High Performance Liquid Chromatography (HPLC). This method can be performed in anaerobic glove bag settings, requires readily available HPLC instrumentation for subsequent detection, and is adaptable to meet the needs of a variety of substrate affinity measurements.

## 1. Introduction

Understanding and quantifying substrate binding is fundamental to decoding enzyme function. When combined with sequence or structural information, determining the *K*_d_ value for a given substrate can accelerate the design of novel biocatalysts and the identification of promising new medically active compounds by allowing researchers to better map structure to function in an enzyme. Currently, there are numerous methods available to measure the binding affinity of a substrate with some popular techniques being isothermal calorimetry (ITC) (Bastos et al., 2023), surface plasmon resonance (SPR) (Olaru et al., 2015), differential scanning fluorimetry (DSF) (Niesen et al., 2007), and fluorescence polarization (FP) (Rossi & Taylor, 2011). The optimal strategy for substrate affinity determination greatly depends on the enzyme, the substrate being probed, the experimental conditions under which the affinity needs to be measured, and the resources and instrumentation readily available.

Non-heme iron halogenases are a family of enzymes capable of directly halogenating an unactivated aliphatic carbon using a mononuclear iron center (Blasiak et al., 2006; Voss et al., 2019; Wilson et al., 2022). Their ability to chlorinate and brominate a variety of substrates including amino acids, nucleotides, and natural products make them a highly versatile biocatalytic platform that invite further optimization. In addition to halogenation, these enzymes have also been shown to functionalize C-H bonds with azide (Büchler et al., 2022; Chan et al., 2022; Matthews et al., 2014; Neugebauer et al., 2019) and nitrite (Matthews et al., 2014), albeit with disappointingly low yields. A variety of methods have been used to determine substrate affinity in these and related classes of enzymes, including tryptophan fluorescence (Martin et al., 2019), SPR (Hu et al., 2015), and UV-Vis spectral shift-based methods (Chan et al., 2022; Matthews et al., 2014; Price et al., 2003; Ryle et al., 1999) to name a few. During our investigations of BesD, a non-heme iron halogenase that natively chlorinates lysine at the γ-position (Marchand et al., 2019), we characterized a positive heterotropic cooperativity that exists between the primary substrate lysine and the co-substrate chloride (Smithwick et al., 2023). To quantify how the concentration of lysine influences the binding of chloride, we were able to utilize a UV-Vis spectral shift-based assay which takes advantage of a perturbation that occurs in a metal-to-ligand charge transfer (MLCT) band between the ferrous iron center and the co-substrate 2-oxoglutarate (2OG) upon the direct binding of chloride to iron (Matthews et al., 2014). In other non-heme iron enzymes, the substrate has also been observed to indirectly perturb this MLCT band by inducing the dissociation of an aqua ligand initially bound to the iron center before substrate binding (Ho et al., 2001). Quantifying this shift upon increasing substrate concentrations has also allowed researchers to determine binding affinities for substrates of non-heme iron enzymes (Price et al., 2003; Ryle et al., 1999). However upon the addition of 5 mM lysine to the Fe^II^-2OG-BesD complex in the absence of chloride, no significant shift in the MLCT was observed. While the absence of any shift could suggest negligible lysine binding at these concentrations, a method that can directly measure lysine binding to the Fe^II^-2OG-BesD complex under anaerobic settings was needed to confirm such a hypothesis. In turn, we developed an equilibrium dialysis assay to directly measure the apparent *K*_d_ of lysine to BesD as a function of total chloride concentrations and is described in this chapter. While the procedure presented here focuses on lysine detection, it is highly adaptable to any amino acid substrate. The assay can be performed anaerobically in a glove bag setting which is a crucial requirement for redox active non-heme iron enzymes, and subsequent HPLC quantification of bound/free lysine fractions can be completed outside the glove bag.

## 2. Materials

All buffers were filtered through 0.22-micron filters prior to application on the HPLC.

### 2.1 Quantitation of ligand and calibration curve

1. High-performance liquid chromatography (HPLC) system with PDA detector
2. C18 column
3. HPLC-grade acetonitrile
4. pH probe
5. 5 mM ammonium acetate solution pH 5.0, pH adjusted using glacial acetic acid (*see* **Note 1**)
6. 30 mM sodium tetraborate pH 10.5
7. 6-Aminoquinolyl-N-hydroxysuccinimidyl Carbamate (AQC)
8. Vortex mixer
9. L-lysine
10. Deactivated clear glass 2 ml vials compatible with autosamplers
11. 300 µL volume polypropylene vials compatible with autosamplers
12. 0.2–10,10–100, and 100–1000 μL pipettes and compatible tips
13. Software to integrate peak area
14. 0.22-micron filters
15. Volumetric glassware

### 2.2 Equilibrium dialysis apparatus setup and initial measurements

1. 0.6 mL low adhesion microcentrifuge tubes
2. Scissors
3. Dialysis membrane 6-8 kDa molecular weight cutoff
4. Parafilm
5. Kimwipes
6. Equipment/materials from section 2.1.

### 2.3 Equilibrium dialysis for determination of substrate affinity in BesD

1. Anaerobic glovebag/glovebox
2. UV-Vis spectrometer
3. 0.5 mL Centricons (centrifugal filter devices) with 10-kDa molecular weight cutoff
4. Centrifuge with cooling capabilities or other refrigeration method
5. Purified non-heme iron halogenase
6. Equipment/materials from sections 2.1. and 2.2.

## 3. Methods

### 3.1 Quantitation of ligand and calibration curve

1. Prepare a series of standard solutions of the ligand you intend to measure. In this study, we studied the affinity of lysine to BesD which will be referred to as the substrate or ligand moving forward. A stock of 30 mM lysine in 50 mM HEPES pH 7.5 was prepared using a 50 mL volumetric flask to adjust to the final volume after pH adjustment. Serial dilutions of 30 mM stock lysine solution were performed in 50 mM HEPES pH 7.5 with pipettes to obtain a set of solutions with concentrations ranging from 2 µM to 300 µM.
2. For analysis of amino acid substrates, prepare the amino acid derivatizing agent solution. Dissolve 3 mgs of AQC in 1 mL of acetonitrile to create a ca. 10 mM AQC solution. Vortex this solution vigorously until all the solid dissolves (*see* **Note 2**). It is best to prepare this solution on the day of analysis as AQC is prone to hydrolysis upon exposure to any H_2_O contamination.
3. Combine 50 μL of one of your standard lysine solutions prepared above with 40 μL of 30 mM sodium tetraborate pH 10.5 in a deactivated glass vial. Vortex to combine before adding the derivatizing agent (*see* **Note 3**). Next add 10 μL of freshly prepared AQC solution directly into the sample. Vortex immediately after each addition (*see* **Note 4**). Repeat for all concentrations of standard lysine solutions.
4. Transfer derivatized samples to 300 µL volume polypropylene vials or other vials that work well with HPLC system of choice. If not immediately analyzing samples, store samples at -20 or -80°C degrees. However, AQC derivatized amino acids have been found to be stable for at least a week (Díaz et al., 1996).
5. Run standard samples on an HPLC to build calibration curve for your ligand of interest. For the analysis of lysine derivatized with AQC, samples were injected onto a Shimadzu Prominence-i LC-2030C 3D Plus system equipped with a PDA detector and a Regis Technologies REXCHROM C18 column (4.60 mm × 250 mm × 5 μm). Solvent A was 5 mM ammonium acetate pH 5.0, and Solvent B was 60% acetonitrile in water. The column temperature was maintained at 35°C. Separation was performed with a gradient method of 0% B to 70% B, 0.0 to 40.0 min; 70% B to 100% B, 40.0 to 42.0 min; 100% B, 42.0 to 50.0 min; 100% B to 0% B, 50.0 to 52.0 min; 0% B, 52.0 to 60.0 min at a flow rate of 0.5 mL/min and an injection volume of 50 μL. All HPLC traces were constructed from extracted absorbance at 254 nm.
6. Build a calibration curve from the ligand specific peak area of the HPLC traces. Peak areas of the three resulting peaks from derivatized lysine products (both mono- and di-derivatized products) were integrated using LabSolutions software (**Fig. 1A**). The total adjusted peak area of all three AQC-tagged lysine peaks were graphed versus concentration of lysine to create a calibration curve (**Fig. 1B**, *see* **Note 5**).

**Figure 1.**
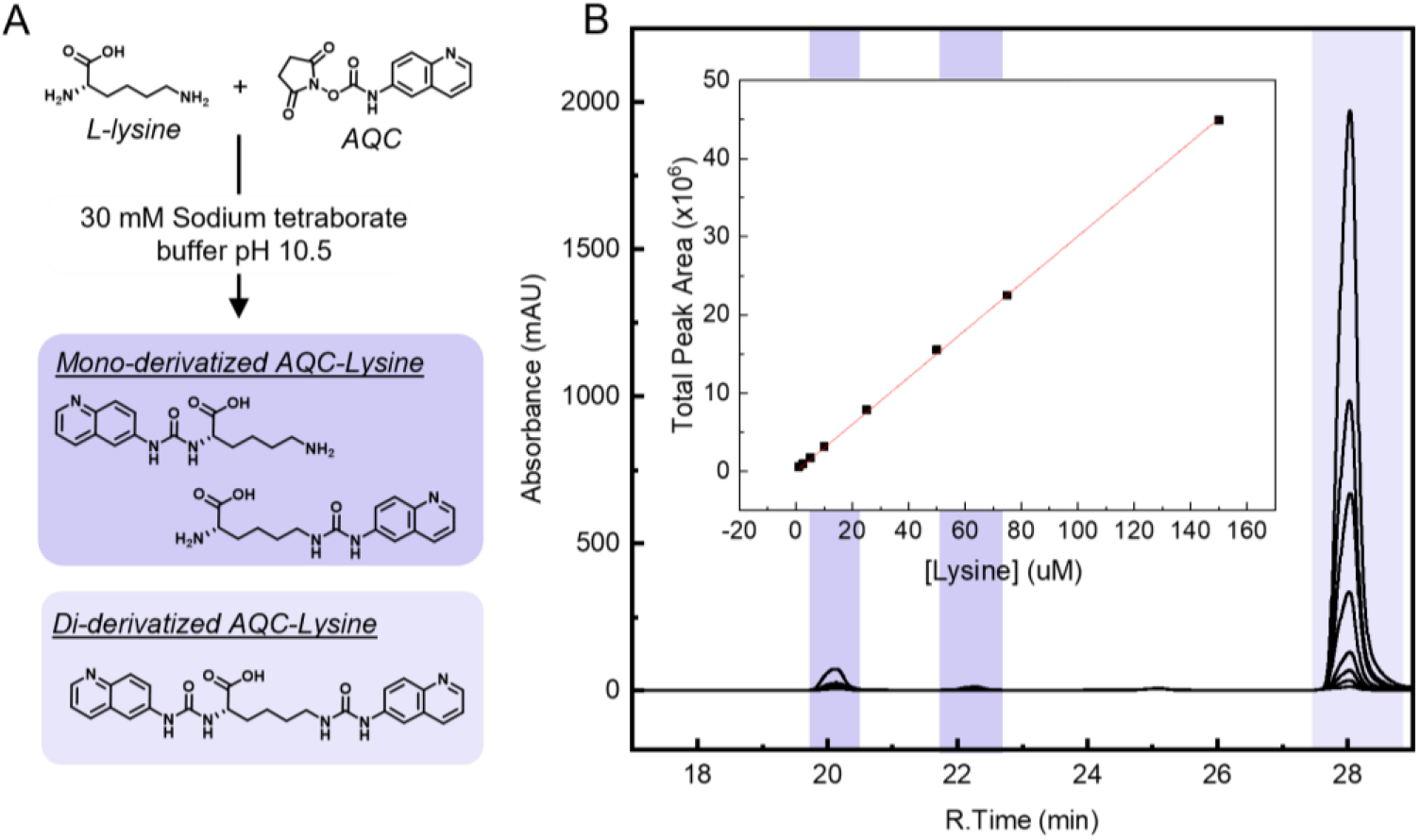
**(A)** Reaction between AQC and lysine produces both mono- and di-derivatized lysine products. **(B)** HPLC traces from AQC-derivatized standard lysine with calibration curve relating relevant peak area to sample concentration shown as an inset. Reprinted with permission from (Smithwick et al., 2023). Copyright 2023 American Chemical Society.

### 3.2 Equilibrium dialysis apparatus setup and initial measurements

1. To prepare equilibrium dialysis setup as previously described (Mega & Hase, 1991), cut a 0.6 mL Eppendorf tube approximately 6 mm below the cap to create a tube open on one side (see **Note 6**). Discard the bottom portion of the Eppendorf tube not connected to the cap. Cut the cap off the top of the tube so the tube portion is completely separated from the cap (**Fig. 2A**). Cut a second cap from a different Eppendorf tube.
2. Cut dialysis membrane into small ~10-15 mm tall pieces and snip the edges of the tubing so that the membrane will be one layer thick. Wash the membrane pieces thoroughly with water and soak them for at least 30 minutes in buffer of choice.
3. Taking one of the previously cut caps, place it face up such that it can be used as the bottom chamber for dialysis (**Fig. 2B**). For initial tests, fill the bottom chamber with 65 μL buffer of choice, in this case, 50 mM HEPES pH 7.5. Retrieve a hydrated dialysis membrane and remove excess liquid on either side with a clean kimwipe. Do not allow dialysis membrane to fully dry out. Gently place the dialysis membrane on top of the bottom chamber. Avoid having any solution that is filled in the bottom chamber from being pulled out by capillary action. Once the membrane is well positioned on top, carefully push the previously cut Eppendorf tube down on the bottom chamber to trap the membrane in between them (**Fig. 2B**). Make sure that the solution in the bottom chamber is in contact with the dialysis membrane at this point. If the solution does not look flush with the membrane very gently tap the tube apparatus until it makes contact. It is common to see a small air bubble off to one side of the tube cap.
4. The trapped dialysis membrane along with tube pressed down upon it forms the floor of top chamber. Add 65 μL of the ligand solution to the top chamber, and then close it up by pushing the second cap onto the open portion of the tube (*see* **Note 7**) (**Fig. 2B**). After the top chamber has been capped, wrap parafilm around the entire apparatus to prevent evaporation of samples. Do not allow samples to tip at this point as liquid getting on sides of the tubing can lead to inaccuracy in quantitation.
5. Allow samples to equilibrate at 4° C. During initial studies, it is best to test a variety of equilibration times to ensure that the highest concentration ligand of interest fully equilibrates between the top and bottom chambers. It was determined that 20 hours was sufficient for complete lysine equilibration at the concentrations of lysine used to evaluate *K*_d_ of BesD.
6. After equilibration, carefully remove parafilm around the equilibrium dialysis setup and then take off the cap of the top chamber. The cap is sometimes difficult to remove so be careful not to remove the bottom cap at the same time as removing the top cap.
7. Transfer 50 μL from the top chamber to a deactivated glass vial and wipe out excess solution with clean kimwipe. Once excess liquid from the top chamber is removed, puncture the dialysis membrane with a piece of wire or a needle and transfer 50 μL from bottom chamber to a different deactivated glass vial.
8. Derivatize all lysine samples with AQC and analyze on HPLC as described in the previous section. Integrate the peaks corresponding to derivatized lysine in each of the sample chromatograms. Using the calibration curve constructed in the previous section, convert integrated peak area to concentration. For samples without protein, fully equilibrated samples should have equal concentrations between the top and bottom chambers and sum to the initial concentration of the solution added to the top chamber. Select an equilibration time for future samples in which the highest concentration of substrate you intend to measure has fully equilibrated between the two chambers. A schematic overview of the equilibrium dialysis process is given in **Fig. 3A**.

**Figure 2.**
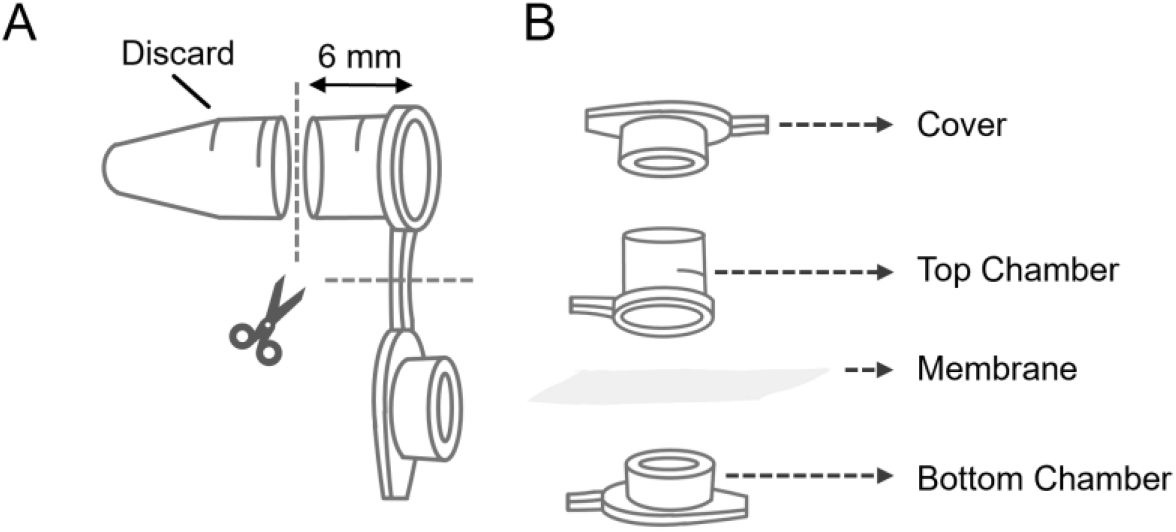
Schematic of the equilibrium dialysis apparatus setup **(A)** Cut centrifuge tube approximately 6 mm from the cap and discard the bottom of the tube. Separate cap from the reserved tube portion and keep both pieces for later. **(B)** Assemble equilibrium dialysis apparatus by adding solution to one of the cap lids placed facing up which will act as the bottom chamber. Position the dialysis membrane in between the bottom chamber and the reserved tubing, and then push down to seal the bottom chamber. Add solution into the resulting open tube portion so that it rests on top of the dialysis membrane. Push a second cap down on top of the open tube to seal the top chamber.

**Figure 3.**
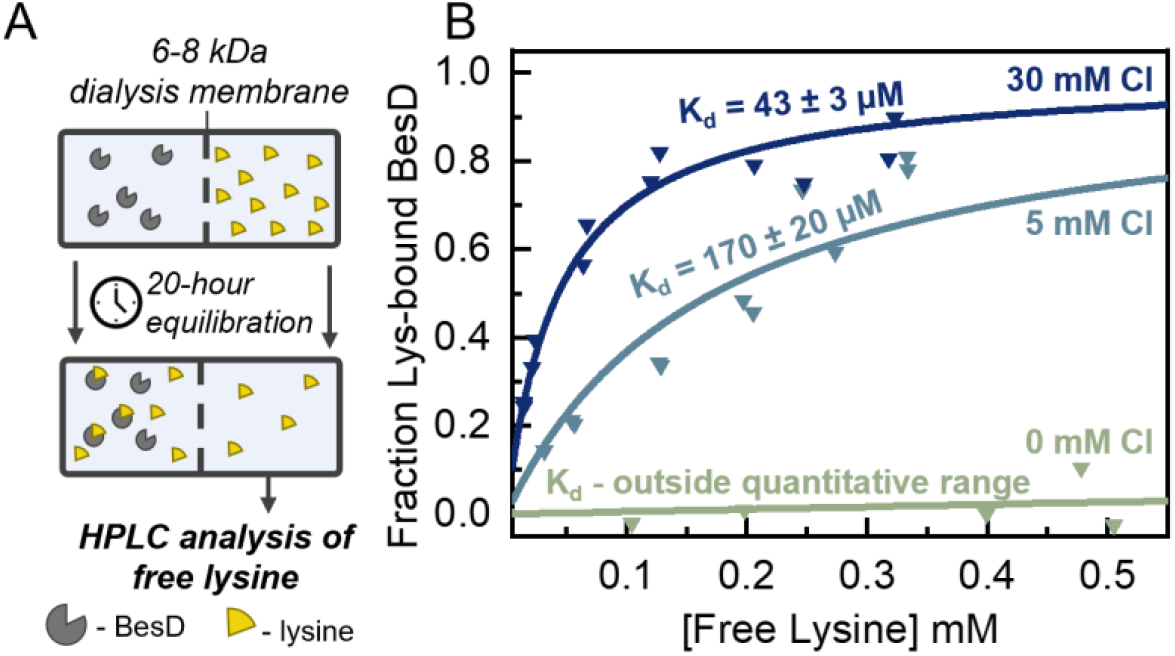
**(A)** Diagram of conceptual equilibrium dialysis workflow. **(B)** Equilibrium dialysis data and derived *K*_d,app_ for lysine binding to BesD at different concentrations of chloride. Reprinted with permission from (Smithwick et al., 2023). Copyright 2023 American Chemical Society.

### 3.3 Equilibrium dialysis for determination of substrate affinity in BesD

1. Bring in materials/chemicals required for equilibrium dialysis experiment into the anaerobic chamber at least 24 hours ahead of the experiment to ensure proper equilibration and removal of oxygen. Deoxygenate any necessary solutions using a Schlenk line, cycling between vacuum and argon gas for 15-minute intervals for a total of three times. Bring into the anaerobic chamber at least 24 hours ahead of the experiment to ensure complete deoxygenation.
2. The day of the experiment, bring in a solution of concentrated stock protein into the anaerobic chamber. Once inside, wash the protein in centricons with deoxygenated buffer by concentrating the protein solution through centrifugation, adding buffer to refill the filter device, and concentrating again. Repeat this process three times to remove oxygen and any other storage buffer molecules. In this study, we examined the substrate affinity of lysine at different concentrations of NaCl in BesD, and our buffer of choice was 50 mM HEPES pH 7.5 for the washing step.
3. After the washing step is complete, invert the filter device in a clean centrifuge tube and centrifuge gently for approximately a minute to reclaim the protein. Assess concentration of protein using absorbance at 280 nm with molar absorptivity calculated by ExPASy ProtParam (Gasteiger, 2003).
4. Prepare the ligand and protein for equilibrium dialysis. In our study, the top and bottom chamber solution consisted of 0.2-1 mM lysine, 4 mM 2OG, 1 mM (NH_4_)_2_Fe(SO_4_)_2_, 30 mM NaCl, and 35 mM Na_2_SO_4_ in 50 mM HEPES (pH 7.5) and of 425±25 μM BesD, 4 mM 2OG, 1 mM (NH_4_)_2_Fe(SO_4_)_2_, 30 mM NaCl, and 35 mM Na_2_SO_4_ in 50 mM HEPES (pH 7.5), respectively. All solutions were prepared from degassed stock solutions of reagents, except for stock solution of (NH_4_)_2_Fe(SO_4_)_2_ which was prepared from dry solid dissolved in degassed MQ water. To assess how the binding affinity of lysine changes with respect to NaCl in BesD, this process was repeated for samples containing 0 mM NaCl with 45 mM Na_2_SO_4_ and 5 mM NaCl with 42.5 mM Na_2_SO_4_ instead of 30 mM NaCl with 35 mM Na_2_SO_4_ while keeping all other concentrations constant.
5. Prepare the equilibrium dialysis setup as described in the above section with the protein solution in the bottom chamber and the ligand solution in the top chamber.
6. After preparation, place sample in refrigerated area set at 4°C. Allow samples to equilibrate for 20 hours. We recommend stirring liquids in the two chambers using stir bar or an orbital shaker, if feasible.
7. After equilibration, recover samples from equilibrium dialysis setup as described in section 3.2. Additionally, take an extra ~5 μL sample of the top chamber and assess if any protein leaked through to the top chamber through increases at 280 nm. While the protein should not be able to traverse the membrane, this check is necessary to verify the integrity of the equilibrium dialysis setup. If protein is found in the top chamber, identify if this is an isolated issue. If the leakage occurs in all samples, consider using a different variety of dialysis membrane (*see* **Note 8**).
8. If no protein leakage has occurred, derivatize extracted top chamber samples and run HPLC analysis using the method described in previous sections.
9. Convert peak areas of sample to concentration using calibration curve to determine the free lysine concentration in each equilibrium dialysis sample. The Hill’s equation for single site binding is given as:

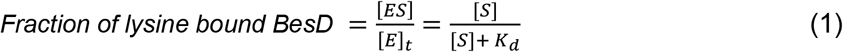

Where [*S*], in this study, is the concentration of free lysine in the top chamber post dialysis as determined by HPLC, [*E*]_*T*_ is the concentration of BesD in the bottom chamber, and [ES] is the concentration of BesD-lysine complex. As [*S*]_*T*_ = [*ES*] + 2[*S*], [*ES*] can be calculated by rearranging the relationship to be [*ES*] = [*S*]_*T*_ − 2[*S*], where [*S*]_*T*_ is the pre-equilibrium concentration of the ligand in the top chamber. Using values of 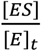 and [*S*], construct a relationship of points and fit to equation (1) using Origin or other software capable of performing non-linear regression:

## Conclusions

Using equilibrium dialysis methods for lysine binding to BesD, we were able to determine apparent *K*_*d*_ for lysine as 43 ± 3 μM and 170 ± 20 μM at chloride concentrations of 30 mM and 5 mM respectively **(Fig 3B**) (Smithwick et al., 2023). In the absence of any chloride, these equilibrium dialysis measurements found negligible binding of lysine to BesD which matched with our observations from UV-Vis spectral shift studies. Overall, the enhanced apparent lysine binding affinity with increasing chloride concentrations measured via equilibrium dialysis approaches in tandem with the enhanced apparent chloride binding affinity with increasing lysine concentrations measured using UV-Vis spectral shift-based approaches demonstrate strong positive heterotropic cooperativity in the binding of lysine and chloride to BesD’s catalytic core. Together they provide insight into the unique ways enzymes can regulate the assembly of their catalytic core and ultimately their biochemical activity. Overall, equilibrium dialysis has the potential to be highly adaptable to a variety of enzyme-substrate pairs and can be easily translated to suit a variety of situations, such as an anerobic environment, due to the low barrier of experimental setup.

### Notes

1. The pH of ammonium acetate buffer solution is very important to reliably measure the retention time of analytes, and deviations of even ±0.1 pH units can affect retention time of the analytes (Van Wandelen & Cohen, 1997).
2. This solution is on the border line of AQC solubility but should be able to completely dissolve in acetonitrile at a ratio of 3 mg/ml. Hold vial in hand to try to warm up the solution to help all the AQC dissolve in solution.
3. We have observed decreased derivatization performance if deactivated glass vials are not used.
4. The pH of the derivatization solution is very important for the AQC derivatization process, so it is important to premix the solutions before adding the AQC. For optimal derivatization, the pH of solution before AQC is added should be between 8.5 and 10.0. When the indicated buffers are combined in the ratios described above, the pH of solution before derivatization is approximately 9.5.
5. Mono-derivatized AQC-lysine products were approximated to have half the absorbance of the di-derivatized AQC-lysine product and therefore peak area corresponding to the monoderivative species were multiplied by 2 when summed to create calibration curve.
6. Instead of the setup we describe here, a variety of pre-made equilibrium dialysis chambers are also available for direct use. For example, Thermo Scientific™ Rapid Equilibrium Dialysis (RED) Inserts and Plates.
7. Depending on how rough the edges are, the cap may be difficult to push down but try to push it so it sits flush with the cut side of the tube. Use a new cap if necessary and be gentle because if you apply force in the wrong way, it will flip the samples and you may have to start over.
8. We did not observe any protein leakage to the top chamber, but this can be an issue if you are working with a smaller molecular weight protein (MW < 20 kDa). Additionally, protein leakage might be evidence that the membrane is being punctured during the assembly process. Make sure to check the integrity of the membrane when removing samples.

## Acknowledgements

ERS acknowledges the support of the National Institute of Health Chemical Biology Training Grant (T32GM132029). This work was supported by NSF CBET and CLP (grant # 2046527).

## Author Contributions

ERS, ABD, and ARD designed the study. ERS performed the study. ERS, ABD, and ARD wrote the article. All authors have given approval to the final version of the manuscript.

